# Continuous mark-release recapture to improve estimates of movement and survival of the African malaria mosquitoes

**DOI:** 10.64898/2026.06.24.734339

**Authors:** M Diallo, A Dao, ZL Sanogo, K Cisse, B Coulibaly, D Samake, BJ Krajacich, A Assitoun, M Traore, J Poudiougo, R Bamou, C Kouam, R Faiman, AS Yaro, T Lehmann

## Abstract

Despite extensive efforts to understand the population biology and ecology of the African malaria mosquitoes questions regarding their movement pattern, survival, and population size persist, reflecting methodological limitations. Site fidelity, in which mosquitoes return to feeding sites, resting sites, or oviposition sites remain debated. Mark release recapture (MRR) studies are vital to address such questions. Using locality- and date-specific DNA tags in fluorescent spray, we carried out a continuous MRR in a Malian village from September to December 2019 with three days interval between capture and release across seven zones. A total of 12,937 *Anopheles gambiae* s.l. (7,455 females) were captured during 35 indoor collections. Handling related mortality was 3.4%., *A. coluzzii* predominated (89.7%), followed by *A. gambiae* (9.4%), and *A. arabiensis* (0.9%). Overall recapture rate was 1.05% (N=129). Contrary to the site-fidelity hypothesis, the distribution of recaptured mosquitoes across zones (regardless of their zone of release) was similar to the distribution of the captured mosquitoes (r=0.97, P<0.001), with 70% recaptured in a different zone. There was no difference in distance moved between sexes, but males average distance increased over time since release, whereas females’ distance remained unchanged. Simulated movements (across released points), with equal probability to reach any of the village houses predicted actual distance moved by mosquitoes. The regression of observed distance from each zone over predicted had a slope of 1 (r^2^=94%, P=0.006), suggesting that the layout of the capture area greatly affected the results. The average days post release (minimum age of wild captured mosquitoes) for recaptures was 6.4 d with the longest being 30 d. No seasonal and sex related difference in minimum age were detected. The corrected probability of daily survival (PDS) was 94% and the daily increase in sporozoite rate was 4.9%. Limiting the recapture duration period showed that PDS increased with recapture duration from 74% to 86% (12 to 30 d, uncorrected). Thus, larger recapture area and longer recapture duration are needed to obtain accurate estimates of movement range and of daily survival.

## INTRODUCTION

Mosquitoes transmit many diseases worldwide, including malaria, yellow fever, and dengue. Malaria’s burden is the heaviest, with 263 million cases and nearly 600,000 deaths in 2024, of which more than 90% occurred in sub-Saharan Africa (WHO 2025). Members of the *Anopheles gambiae* s.l. complex are the primary malaria vectors in Africa. Effective vector control requires knowledge of the natural history and ecology of the mosquito vector. Despite a century-long research, vector components of disease transmission by the African malaria mosquitoes remain poorly understood (Chitnis et al. 2008, Brady et al. 2016). Mark-Release-Recapture (MRR) studies represent the ‘gold standard’ method to estimate survival, movement distance range, and population size of field mosquitoes (Gillies 1961, Guerra et al. 2014). However, inconsistent estimates of survival (expected life span), movement range (dispersal), and population size (Service 1993, Faiman et al. 2022) are common, reflecting methodological limitations (Gillies and Wilkes 1965, Guerra et al. 2014, Matthews et al. 2020, Matthews 2025). For example, estimates of daily survival span the range from 0.3 to 0.98 (Matthews et al. 2020, Matthews 2025), with the lower estimates obtained mostly by MRR compared with parasitological and the parity or ovariole dilatation methods. This difference is attributed primarily to confounding mortality with emigration outside the trapping area. Guerra et al. (2014) points to such inconsistencies and stressed that studies with a larger trapping area yielded larger estimates of the mean distance traveled, indicating that most if not all studies have not designed sufficient area size to recapture released mosquitoes, thus underestimating dispersal (mean distance traveled) and survival. Adding to this confusion, is the concept of “site fidelity” in malaria mosquitoes (Charlwood et al. 1988, McCall et al. 2001, McCall and Kelly 2002), albeit controversial (Alonso and Schuck-Paim 2006) that implies that mosquitoes tend to return to the same sites to blood-feed, rest, and lay eggs where closer alternative sites are available, thus reducing net displacement and mean distance travelled and especially, limiting pathogen spread.

The typically low recapture rate <3% in anopheline studies (Guerra et al. 2014) and the considerable effort and cost associated with MRR experiments, have led to “economical” design of the experiments with i) small (<800 m radius) trapping area, ii) short duration (<12 d) of recapture, and iii) early first recapture (with daily interval thereafter), i.e., in the day of the release or the following day, allowing insufficient time for mosquitoes to move outside their release area. Compounded by a limited number of distinct tags, i.e., fluorescent colors that are easy to distinguish from each other because many colors are mixes of basic colors that can only be distinguished in high density. Thus, the information produced has been limited and likely biased (Guerra et al. 2014, Matthews et al. 2020, Matthews 2025). For example, different studies estimates of expected adult longevity ranged from 3.6 to 20 days (Matthews et al. 2020, Matthews 2025)

To address some of these issues and evaluate the effect on the estimates of daily survival and movement range with expectations based on previous studies, we have conducted a MRR study in the Sahelian village Thierola over 101 recapture days period (from September 3 to December 13, 2019) and allowed 3 days interval between (multiple) release and recaptures to ensure mosquitoes could complete their gonotrophic cycle (Gillies and Wilkes 1965, Toure et al. 1998). We used the DNA tags mixed with fluorescent solution recently described (Faiman et al. 2021a), to identify mosquitoes released up to 36 days post release. Our results indicate that long recapture period produce longer expected adult longevity (higher daily survival). Additionally, we find evidence for extensive movement at the village scale which implies low or no site fidelity, despite marked heterogeneity in concentration of mosquitoes in the village, both captured and recaptured. These results suggest that to obtain accurate estimates of survival and movement (among other parameters) from MRR studies, longer period of recapture and larger recapture area are essential.

## METHODS

### Study area

The study was performed in the Sahelian village Thierola (13.40° N, 7.13° W) the rural commune of Kiban, Koulikoro region which was previously described in detail (Lehmann et al. 2010, Dao et al. 2014, Lehmann et al. 2014). Briefly, Thierola is surrounded by farmland growing millet, sorghum, and vegetables during the wet season (June to October). The cumulative rainfall in 2019 this area received was 653 mm^3^ with a mean temperature of 27.9°C (based on a weather station set in Thierola). Larval habitats are widespread between July and September but become sparse afterwards. The last larval site within 5 km radius around the village dried up in November. Theirola is located 3 km from the closest village, Zanga and 5-8 km from next neighboring villages. The population is estimated at 500 inhabitants living in 162 houses (Figure 1).

**Figure 1.**
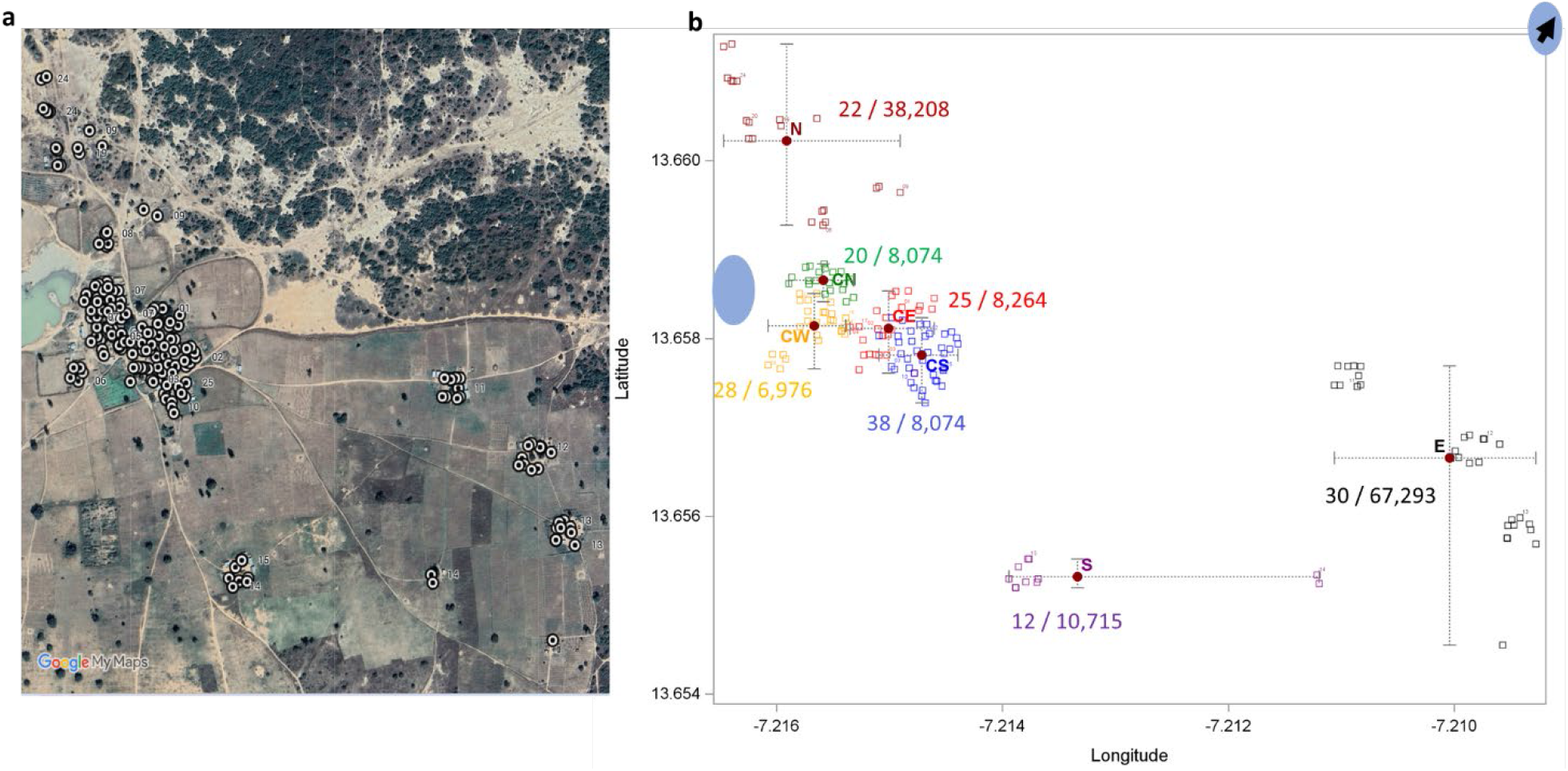
Layout of houses in the study village and division to zones. A satellite image of the village showing houses (circles) and the main larval site (blue ellipse a). The houses in each zones (colors and letter code), the number of houses per zone and its approximated area based on its latitudinal and longitudinal lengths as shown (b). The release point of each zone is shown (filled circle). The position of the second larval site is 1.5 km in the direction of arrow (top right corner).

The village was subdivided into seven zones based on their spatial organization (Figure 1). The number of houses and fraction of houses in each zone are as follows center west (CW): 27 (17%), center south (CS): 37 (23%), center north (CN):19 (12%), center east (CE): 24 (15%), east (E): 29 (18%), south (S): 11 (7%), and north (N): 15 (13%). The largest and more stable larval sites were located ∼25 m from the CN (Center North) and CW (Center West) zones (Figure 1). These larval sites were especially important during the end of the wet season (October-November). Animal enclosures are also present near many households with several large cattle, goat, and sheep enclosures located ∼150 m (or more) from houses of zones E and S.

### Study design

The MRR experiment was carried out continuously from September 3 to December 13. Mosquitoes were collected across the village Thierola (>90% house coverage) every 3^rd^ day. In each day of collection, unmarked mosquitoes were marked following capture and released at sunset (of the same day). Mosquitoes were marked with a color-specific visual fluorescent dye that corresponded to their collection zone (and release point) as well a synthetic DNA tag that matched their zone of collection (and release) and a second DNA tag that marked the date of their release. The date tags were reused every 39 days. Accordingly, mosquito date of release was assumed to be up to 38 days before its recapture date. The interval of three days matches the typical length of *A. gambiae* s.l. female gonotrophic cycle (Gillies and Wilkes 1965, Toure et al. 1998). Different types of movements that occur during a gonotrophic cycle, e.g., host and sugar source location, resting sites location, and oviposition site location, which differ in expected displacement distance and likely in direction are summed over each 3-d, gonotrophic cycle. The net sums of these movements are expected to be more similar to each other than day-to-day distances. Avoiding capture the two days following release prevents capture of mosquitoes that were damaged by handling, which tend to inflate the recapture totals especially near the release site and allows for more homogenous mixing in space. Given the twist and turn of appetitive movements to and fro larval sites, it is the net distance over the gonotrophic cycle that best measures the displacement of *Plasmodium* from host to host.

### Mosquito collection and processing

Mosquitoes were collected indoors between 08:00 and 10:00 AM using a mouth aspirator and flashlight. Each house was thoroughly searched for 5 minutes by two trained collectors. The mosquitoes were gently aspirated into black collection paper cups (height cm x diameter cm), provided with 10% sugar water. The cups were placed in a cool shaded area and covered by a wet cloth to maintain high relative humidity. Each zone had a dedicated room for preparation and marking (below). Mosquitoes collected from each zone were transported to a zone-specific field laboratory, where marked individuals were identified under a UV flashlight inside black containers in a dark room and preserved over silica gel. Unmarked mosquitoes were marked and released at sunset (18:45–19:00) the same day of collection. Additionally, surveys for available larval sites were carried out around Thierola and in closest villages Zanga and Bako, 3 and 5 km away, respectively, where presence/absence of water and anopheles larvae was recorded.

### Mosquito marking

Wild adult mosquitoes were marked using a fluorescent dye SmartWater® with synthetic DNA tags [, applied via an airbrush and nebulizer as described previously [11]. Tag mixture solutions were prepared hours prior to marking. A fluorescent tag solution contained two specific DNA tags, one to indicate the date and the other to indicate the locality of capture and release. The “A series tags” was utilized to determine the dates of capture marking and release. This series consisted of unique fragments: 80, 100, 120, 140, 160, 180, 200, 220, 240, 260, 280, 300, 320, 340 base pairs (bp), allowing distinguishing between 14 unique dates spread through 36 days before repeating on the 39^th^ day from the first release. The “D series” was used as a marker of the location (zone) of capture and release. This series consisted of unique fragments of 70, 90, 110, 170, 190 bp. Each series was amplified using distinct DNA primers [11] and in case of length ambiguity, fragments could be distinguished based on their sequence [11]. In addition to the series D tags, we used four fluorescent colors (yellow, light blue, orange and red) to represent the zones of capture and release throughout the experiment. After marking mosquitoes were placed in groups of 25 per standard cup (7 cm diameter * 9 cm height) for release that evening (below).

### Release process

The releases took place at sunset 18:45–19:00. Marked mosquitoes were taken to their zone-specific release point and allowed to settle for ten minutes before the netting over the top of the cup was removed, and mosquitoes started to fly out. If mosquitoes remain in the cup, we gently tapped on the bottom of the cup. Mosquitoes that were damaged by the procedure and did not fly, were counted and preserved using silica gel.

### Laboratory and Molecular analysis

Following collection, recaptured mosquitoes (and those damaged by handling) were sorted to morphological species and members of the *A. gambiae* complex stored in Eppendorf tubes containing silica gel. Silica gel was chosen due to its effectiveness in preservation of DNA even at room temperatures. Recaptured mosquitoes (previously marked) were reexamined for color determination using a UV torch in dark room. The legs of male mosquitoes were used for DNA extraction for DNA tag determination (below). The legs and head-thoraxes body parts of female mosquitoes were used individually for DNA extraction. DNA extraction was performed using a Trizol-based reagent (TRI Reagent®, Zymo Research, US) in combination with the Mag-Bind® Viral DNA/RNA 96 Kit (Zymo Research, US), utilizing the KingFisher® Flex automated extraction robot (ThermoFisher, US) to ensure consistency in sample preparation. This procedure was previously described in detail (Faiman et al. 2020, Bamou et al 2026). The extracted DNA of individual mosquitoes was used for i) species identification, (ii) screening for malaria parasites (females only), and (iii) detection and identification of the DNA tag used for marking mosquitoes. Identification of *A. gambiae* s.l. members was achieved following Wilkins et al. (2006), which distinguishes species based on amplified DNA band sizes observed after gel electrophoresis. Positive controls consisted of *A. gambiae* s.s. and *A. coluzzii* from laboratory colonies, while negative controls included Culex mosquitoes and water. For malaria infection screening, DNA from the head-thorax of female mosquitoes was analyzed using TaqMan SNP genotyping, which enables identification *Plasmoidum falciparum, P. vivax, P. ovale*, and *P. malariae* (Bass et al. 2008). All runs include both positive controls (*P. falciparum* D7 and *P. vivax* from laboratory infected mosquitoes) and water for negative controls.

For the identification of the DNA tags used for mosquito marking, conventional PCR followed by gel electrophoresis was performed separately for the A and D marker series (above). The PCR reaction consisted of 6.25 μL of Ready-to-Go Master Mix G2 Gotaq (Promega, US), 0.125 μL each of forward and reverse primers (100 μM) specific to the series (A or D), 0.15 μL of MgCl2 (25 mM stock), 4.85 μL of DNA-free water and 1ul of genomic material for a final volume of 12.5ul. For each PCR run, positive controls included corresponding DNA tags (A or D), and negative controls comprised unmarked mosquitoes from the field and laboratory, as well as a no-template (water) control. Amplicons were loaded on 2% gel electrophoresis for band size verification.

### Data analysis

All data were recorded in Excel and analyzed in SAS (SAS software 2019). To understand variation in collection density across the village over time we compared Poisson and binomial regression models with autoregressive errors implemented in PROC AUTOREG and GLIMIX (SAS software 2019). Chi-square test was used to compare mortality rates (proportions) of males and females and to compare simple recapture probabilities given those released. Pearson correlation coefficients were computed using PROC CORR and 95% confidence interval of the means (CI) were generated as part of the statistical graphics procedures of SAS, e.g., PROC SGPLOT (SAS software 2019).

To evaluate if mosquito movement distances was affected by the layout of the experiment and village, we simulated 1,000 replicated recapture samples, each of 126 “mosquitoes” stratified by the zone of release, so that the same numbers actually released and recaptured from each zone were equally represented in each simulation, but that all houses had equal chance to be selected by a mosquito (sampling with replacement). These simulations were performed using PROC SURVEYSELECT in SAS (SAS software 2019). The mean distances of the simulated samples were compared with the actual mosquitoes.

Daily survival rate or probability of daily survival (PDS) was estimated following Gillies (1961), by regressing log10 (recaptures+1) over the day of recapture, and computing the antilog(base 10) of the slope of the regression. The average life expectancy (ALE_G_) was computed as 1/-ln(PDS_G_). These estimates are comparable to many others. Additionally, we applied the linear correction suggested by Buonaccorsi et al. (2003) that adjusts for previously recaptured mosquitoes: PDS_GB_ = e^b^ / (1-ϴ)^1/d^, with b representing the slope of the regression above, d reflects the number of recapture days, and ϴ representing the daily recapture rate rather than the simple recapture rate (total recaptures/total released) over multiple days of capture: ϴ = e^a^ / (N+ e^a^), with a representing the intercept of the regression above, and N representing to total number of released mosquitoes.

## RESULTS

From September 2 to December 13, 2019, a total of 12,937 *A. gambiae* s.l. (7,455 females, 57.6%) were captured during 35 indoor collections 3 d apart. The collection coverage amounted to 90.2% of all house-structures (N=**162**, including unoccupied houses, kitchens, storage, etc.). Mosquito density indoors varied over time and across zones (**Figs. 1 and 2**). As expected, (Dao et al. 2014), during this transition time (wet to the dry season), indoor density of *A. gambiae* s.l. has declined sharply from its peak in mid-September to early November and then declined slowly until the end of the experiment in its mid-December (Fig. 2b). We also estimate the change in relative indoor density per each time point (Fig. 2b: inset) based on the local regression predicted values of house density (D) at each consecutive time points as: 1 + [(D_t_ -D _t-1_) / (time _t_ -time_t-1_)]. It shows a rapid decline in daily growth rate after September 5^th^ (peak) from 1.4 to 0.46 (minimum) on September 23, crossing 1 (stable population growth) around September 15^th^. Subsequently, it slowly grows towards 1 where it hovers from November 4^th^ to December 13^th^ (end of the experiment). The last larval site with 5 km radius from the village dried up on November 17, 2019 but its surface and apparently productivity declined from early September. Indoors density varied among zones (Fig. 2a) with highest overall density in zone CN (4.13) and CW (3.95) compared with the remaining zones (0.96–2.29, Fig. 2a). Indoors density of females and males were highly correlated (r=0.91, **Fig. S1**). Indoor mosquito density per zone was correlated with the zone’s house density but not with its area or number of houses (Figs. 2c, 2d, and S1c). Although the species composition among the mosquitoes released is unknown, using mosquitoes that died during processing or did not fly during release (below), the overall composition of *A. coluzzii, An, gambiae s*.s. and *A. arabiensis* were 89.7%, 9.4%, and 0.9%, respectively (N=435).

**Figure 2.**
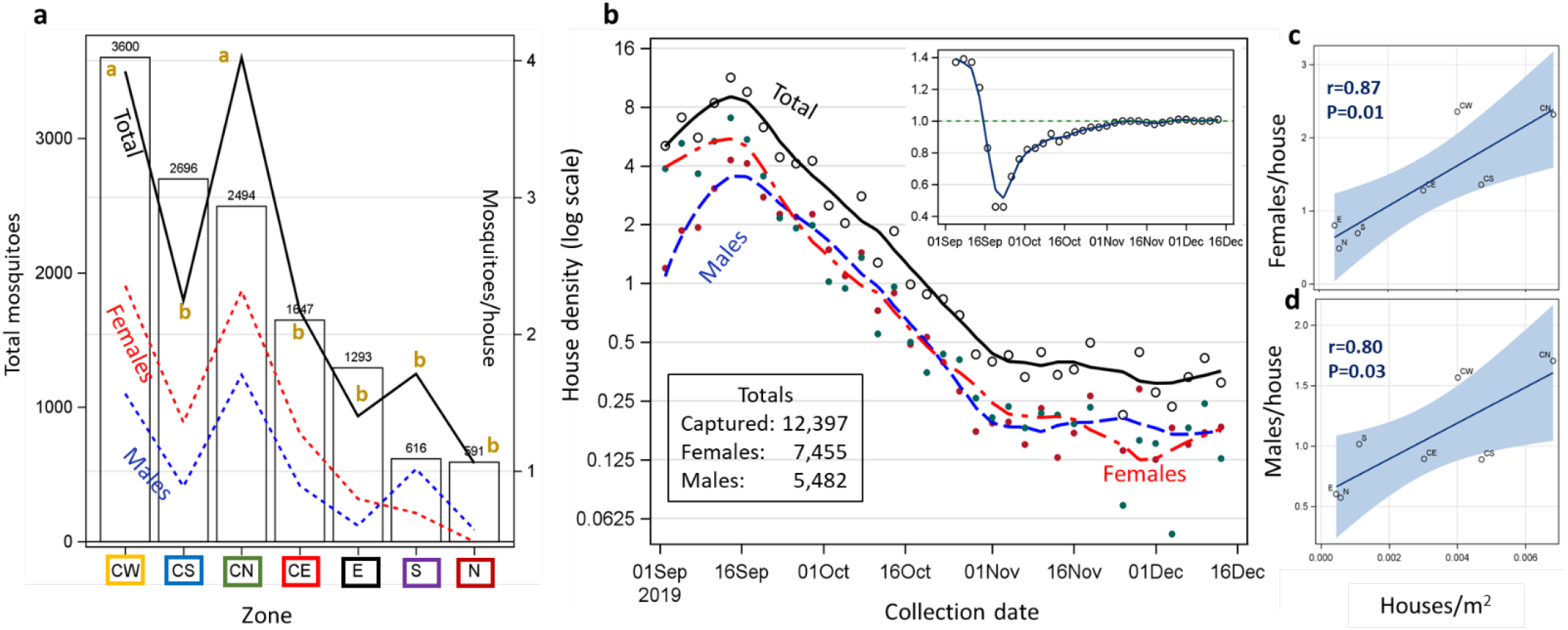
Abundance of *Anopheles gambiae* s.l. during the experiment by zone (a), season (b) and in relationship to house density per zone area (c). Total number of *A. gambiae* s.l. captured by zone (bars) and corresponding mean indoor density (lines) over the study period. Zone colors correspond to those in Figure 1 (**a**). Dynamics of indoor density of total, female and male mosquitoes using local regression (loess) over the experiment (**b**). Inset: population growth estimate (λ) measuring change in density (D_t_ -D _t-1_ / time _t_ -time_t-1_). The relationships between indoor density in each zone and house density are shown for females and males (**c and d**) with a regression line and 95% CI and the associated Pearson correlation coefficient (N=7 zones).

Mortality rate during the processing of 12,937 mosquitoes (capture-mark-release) was 3.4% (N=436). Mortality was higher in males (4.1% N=5,773) than in females (2.8%, N=7,758, χ^2^_df=1_ = 16.5, P=0.0001). There is no information on gonotrophic stage of female released so comparisons reflect the proportions of gonotrophic states among those that died: gravid (36%), half gravid (33%), blood-fed (19%), and unfed (12%). Species composition was dominated by *A. coluzzii* (89.7%), followed by *A. gambiae* (9.4%) and *A. arabiensis* (0.9%). The species composition was not different among males and females (N=435, χ^2^_df=1_ = 1.7, P=0.4). Overall sprozoite infection rate with *P. falciparum* in head-thorax was 3.3% (7 of 210 females) and overall anthropophagy was 96.3% (N=108 blood fed females). No significant differences were found between species in infection rate or anthropophagy (χ^2^ _df=1_<0.98, P>0.6).

Overall, recapture rate was 1.05% (129 recaptured of 12,343 released mosquitoes, **Fig. 3**). The capture and release zone was determined for 128 and date of release was determined for 129. The species composition among recaptures was 90% *A. coluzzii*, 8% *A. gambiae*, and 2% *A. arabiensis* (N=104). No recaptures were found from mosquitoes collected and released at zones CE and CN, despite robust numbers of releases mosquitoes and the lowest mortality rates (Fig. 3a). The overall distribution of recaptured mosquitoes across zones (regardless of their zone of release) was very similar to the distribution of the captured mosquitoes (Fig. 3b). This could suggest complete mixing of released mosquitoes in the village or that the vast majority of released mosquitoes remained in their release zone (zone fidelity). Considering the capture and release zone of the mosquitoes, 29.7% have been recaptured in the same capture zone, whereas 70.3% have been recaptured in a different zone (Fig. 3c).

**Figure 3.**
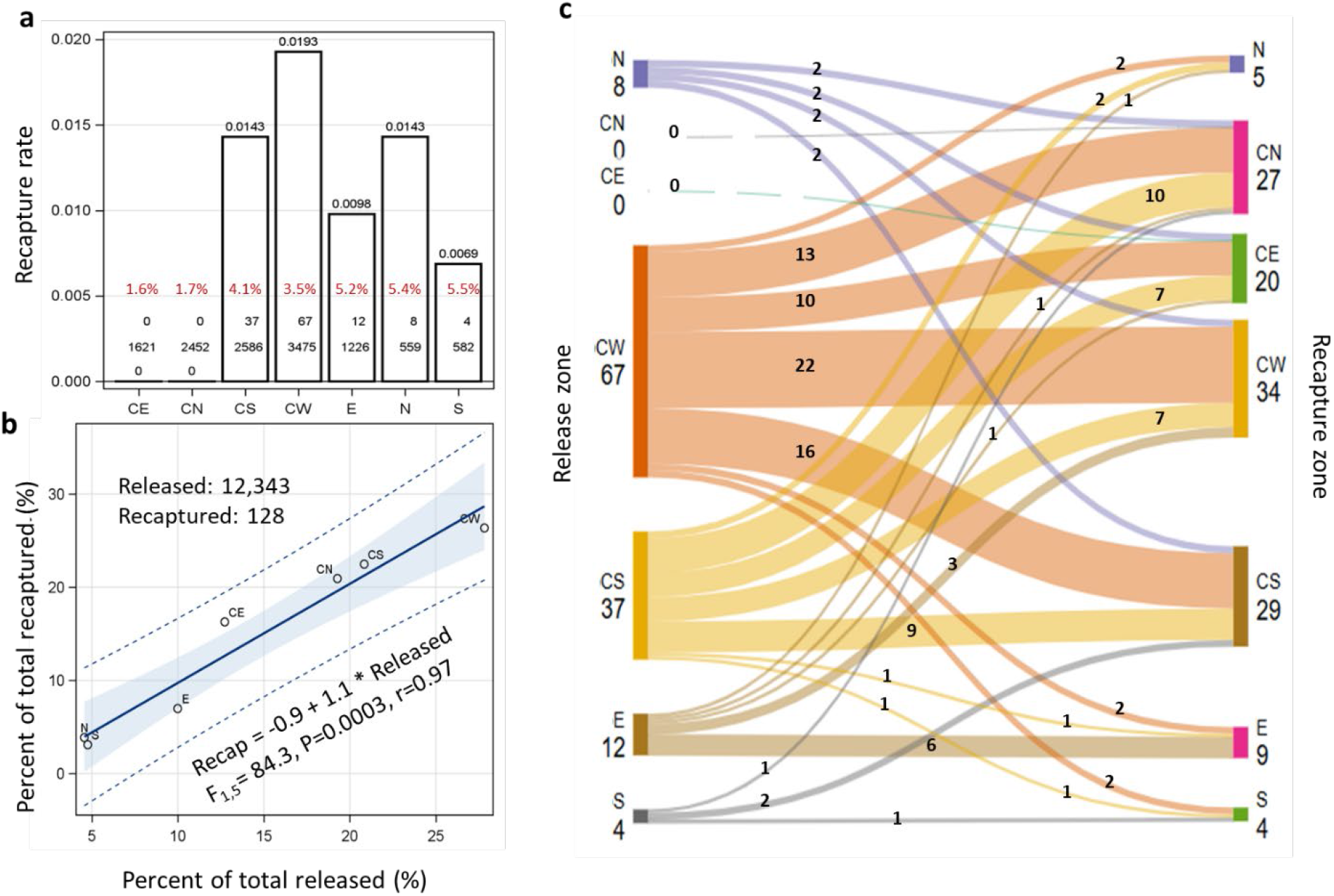
Recapture rates and distribution of recaptures across zones. Zone-specific rates of recaptures showing totals released and recaptured from each zone at the base of the bars as well as mortality rate in red (**a**). The relationship between the percentage of mosquito captured in each zone and the percentage of recaptured mosquitoes across zones (regardless where they were released from) with regression line (blue) surrounded by 95% CI from the mean (band) and 95% CI for individual observation (dotted line; **b**). Sankey diagram showing distribution of recaptured mosquitoes with their zone of release and recapture (**c**). Line thickness is proportional to number of mosquitoes (black numerals). Thin broken lines from CN and CE represent zero recaptured mosquitoes from these zones.

### Distances moved and experimental layout

The mean distance moved by recaptured mosquitoes was 136 m (max= 623 m, Fig. 4a). There was no sex difference (males: N= 80, mean= 149 m, max= 566 m; and females: N= 46, mean = 112 m, max= 623 m, Fig. 4a). Both male and female mosquitoes covered nearly the same maximal distance in 3 days (one gonotrophic cycle), but fewer males remained near the release sites over time, producing a trend of increased distance over time (days since release, Figure 4b). Females exhibited no such trend (Fig. 4b). Notably, the maximum possible distance a mosquito released in this experiment could reach is 886 m, between the release point at the northern zone (N) and a single remote house south of zone E (Fig 1). Because only 8 mosquitoes that were released from this zone were recaptured, the likelihood that one will reach that house is rather small (∼4.8% = 8/168). To determine the degree in which the layout of the experiment (and the village) shaped the scope of mosquito movement, we used Monte Carlo simulations generating 1,000 replicates samples of 126 recaptured mosquitoes, stratified by their released point, with equal probability to reach each one of the 168 village houses (Methods). We then compared the distance distributions of these simulated “releases” with the actual sample (Fig 4c). The simulated distance (230.4 m) was 1.7 times greater than that of the recaptured mosquitoes (136 m, Fig. 4c). The regression of actual distance moved by mosquitoes from each zone on their corresponding simulated distance was highly significant (P=0.006), with a slope of 1 and intercept of -102 m. The r^2^ coefficient was 94%, demonstrating that a large proportion of the variation in recapture distances is accounted by the experimental (including the village) layout. Note that movement increased among zones with distance from the village center, which is also near the main larval site (Fig. 1b and Fig 4d).

**Figure 4.**
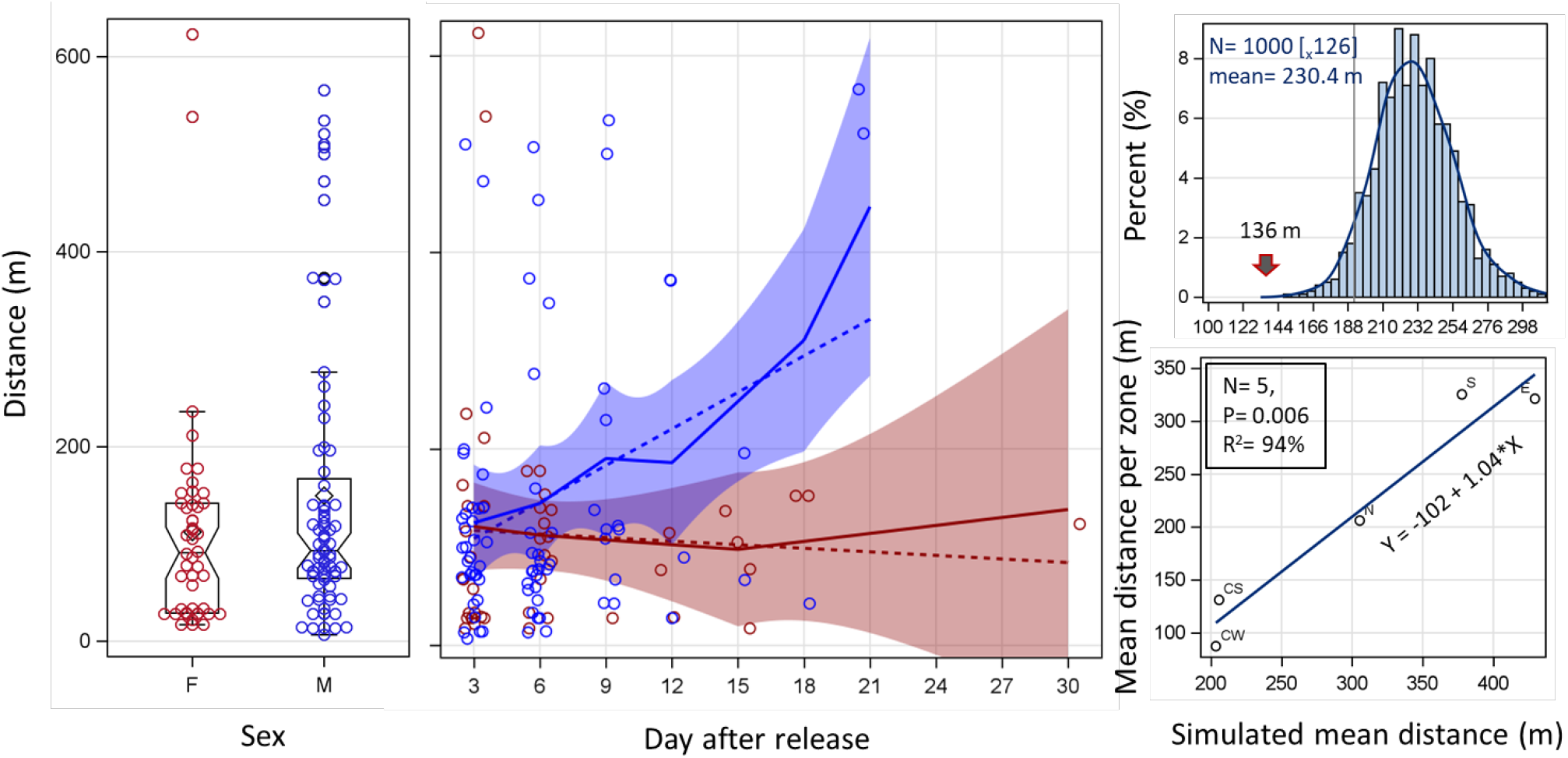
Linear distance between release and recapture points of males and females (a and b) and the difference between empirical and randomized distance distributions (c and d). Distributions of linear movement distance by sex (a) using box whisker plots, with upper and lower box sides showing the first and third quartile, respectively. The median and mean are shown as horizontal lines and diamond. The whiskers extend up to 1.5 times the inter-quantile range (Q3-Q1) and putative outlier values are above the whisker ends. The effect of time until recapture on linear distance (b) for males (blue) and females (red) with local (continuous) regression with 95% CI and and linear regression (broken line) for comparison. Mean distance distribution of 1,000 simulated movements with equal chance to reach each house (c, see text) with actual mean distance moved (arrow). Regression of mean distance moved by mosquitoes released from each zone on the corresponding simulated distance (d). Regression statistics and equation are shown with zone labels.

### Daily survival estimation

Although our experiment lasted 101 days, we reused the same tags after 36 days, limiting our ability to detect mosquitoes that were recaptured over longer period (Methods). One mosquito was captured 30 d after release and two were caught after 21 d (Fig. 5). The time between recapture and release was 6.4 d, estimating the minimum age of the mosquitoes, revealed no significant difference over time (September to December), release zone, or sex (P>0.07, not shown), despite an apparent trend suggesting that females survived longer than males (Figure 5b, 5c). Covariance analysis to compare the slopes of the log counts of males and females over time since release, detected no significant interaction (F_1/10_=3.9, P=0.076) and no significant main effect of sex (F_1/10_=4.35, P=0.064). Given that the log of the recaptured mosquitoes over time since release remained close to linear up to the last recaptured mosquitoes (30 d, Fig. 5d), we estimated the probability of daily survival (PDS) based on the classic method of Gillies (Gillies 1961) and also based on the corrected method (Buonaccorsi et al. 2003) as 0.86 and 0.94, respectively. To assess the effect of the duration of the recapture period on the estimate of PDS (following Gillies), we sequentially extended the duration from 12 day to 30 days (Fig. 4d: inset). Consequently, the estimate of PDS increased from 78% to 86%.

**Figure 5.**
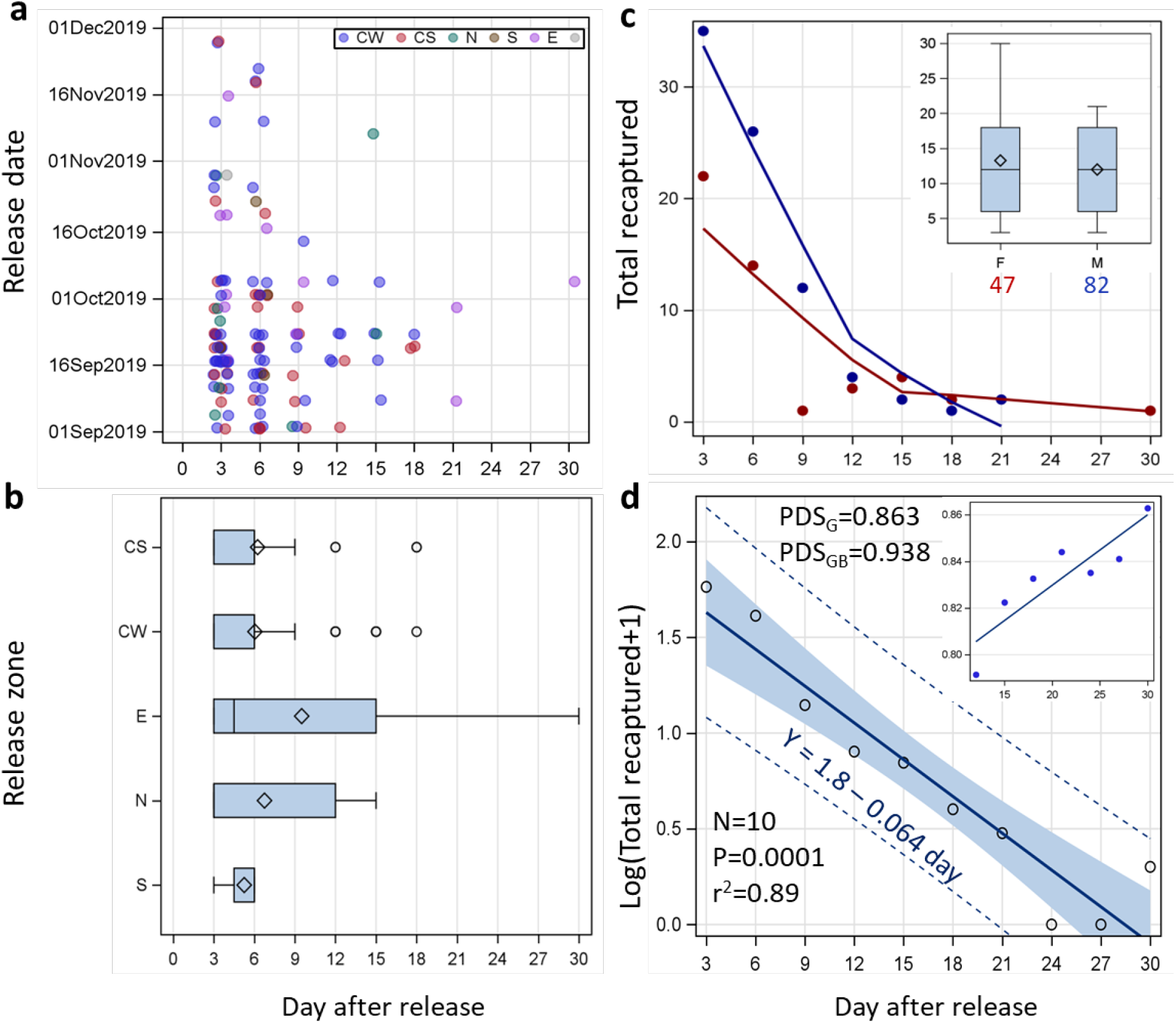
Minimum age of recaptured mosquitoes measured by the number of days between dates of recapture and release and estimate of probability of daily survival (PDS). Distribution of minimum age in relation to date and zone of release (a). Distribution of minimum age in relationship to zone of release (b). Sex specific difference in minimum age (c). Estimation of daily survival (PDS_G_) based on the regression of log(recaptures+1) over days since release (females and males pooled; see text) and the corrected estimate (PDS_GB_). The 95% CI for the regression mean (shaded) and for individual observations (dotted lines) over days and corresponding ALE. Inset: the effect of increasing duration of recapture post release (from 12 to 30 day), on the estimate of daily survival (PDS_G_, see text).

Compared with captured mosquitoes whose overall infection rate with sporozoites (infection in thorax-head, above) was 3.3 %, that in recaptured mosquitoes was 23.9% (N=46). The sporozoite rate increased 15% per gonotrophic cycle (3 d) with time after release based on a regression that included the rate at capture as time zero (5% per day, Figure S2). In that regression, we pooled recaptured mosquitoes collected 9 days or later into one group (N=10).

## DISCUSSION

Knowledge of vector population size, movement patterns, and survival (and their variation) is fundamental to understanding vector-borne disease transmission (Chitnis et al. 2008). It is also essential for planning effective vector and disease control interventions (Brady et al. 2016). MRR studies represent the gold standard of measuring survival, movement range, and population size of wild populations (Reisen et al. 1991, Service 1993, Costantini et al. 1996, Toure et al. 1998, Guerra et al. 2014, Verdonschot and Besse-Lototskaya 2014, Faiman et al. 2022). However, estimates of these fundamental parameters for mosquitoes are often spread out over suspiciously large range, questioning their accuracy and limiting their utility (Guerra et al. 2014, Verdonschot and Besse-Lototskaya 2014, Matthews et al. 2020). Striving to improve the inference on these parameters of malaria mosquitoes, we carried out one of the longest MRR study (101 days) at the Sahelian village Thierola, using a new marking approach (Faiman et al. 2021b) that allows identification of large numbers of subpopulations by locality- and date-specific DNA tags mixed in fluorescent solution, we simultaneously released mosquitoes from 7 zones within the village. A total of 12,937 *A. gambiae* s.l. (90% *A. coluzzii*) were captured, marked, and released during 35 indoor collections, 3 d apart, and 129 mosquitoes were recaptured (1.05%). The species composition among the recaptures was similar, suggesting that there results pertain mostly to *A. coluzzii*. Our results reveal that a longer duration of recapture period improves estimation of survival (expected longevity) of wild mosquito population. We discuss key results and lessons for future MRR studies.

Over the course of the experiment density of *A. gambiae* s.l. declined sharply from early September to end of October (Fig. 1), although recruitment of emergent (young) mosquitoes continued until November 17, 2019 when all larval sites surrounding the village up to 5 kilometers away dried up. This dramatic change highlights that our MRR results primarily refer to September October as the number released and recaptured later were considerably smaller. Afterwards, as density decreased, the rate of population decline increased and reached almost one (stable population, Fig. 1). This low but stable populations density (0.3 mosquito/house, Fig. 1) may reflect immigrants arriving from larval sites 5 km aways that persist until January (Bako), or even from distant rice irrigation fields ∼150 km away (Dao et al. 2014, Faiman et al. 2022), or aestivating adults (Omer and Cloudsley-Thompson 1970, Lehmann et al. 2010, Adamou et al. 2011, Dao et al. 2014, Faiman et al. 2022). Additionally, density varied considerably among zones (Fig. 2a) with highest overall density in centers zones CN and CW (Fig. 2c). Indoor density was correlated with the zone’s house density (Figs. 2c, 2d, and S1c) but also with the location of the main larval sites nearest the village during the wet season (Fig. 1), an association that is widely recognized (Smith et al. 1995, Ribeiro et al. 1996, Carter et al. 2000, Kamdem et al. 2012). This pattern is highly consistent with that described in Thierola during the wet seasons between 2009-2013 (Lehmann et al. 2014).

Mosquito site fidelity refers primarily to mosquito movements between the same resting sites, blood-feeding sites (e.g., houses) and/or larval sites where closer alternative sites are available (McCall et al. 2001). The implications of site fidelity in reducing the spread of pathogens and mosquito genes (e.g., insecticide resistance) are significant, (Charlwood et al. 1988, McCall and Kelly 2002), however, the evidence backing it is inconclusive (Alonso and Schuck-Paim 2006). Because site fidelity reduces the net displacement of mosquitoes, it can be assessed at the village scale. Our results reveal mixing of recapture mosquitoes across zones, contrary to the site-fidelity hypothesis (Fig. 3c). The distribution of recaptured mosquitoes among zones followed closely zone density (capture density, Fig. 3b, r=0.97 P<0.001), with 70% recaptured in a different zone than their capture zone, indicating extensive exchange among zones yielding thoroughly intermingled recaptures over the village (Fig. 3c).

No significant difference was found in mean displacement of females and males (Fig. 4a). Females exhibited rapid mixing in space within one gonotrophic cycle with stable displacement over time since release (Fig. 4b). Males also exhibited rapid mixing within 3 d post release, but their average displacement increased over time (Fig. 4b), as a result of fewer recaptured males near their release sites (rather than larger displacement distances, Fig. 4b), likely reflecting the limitation of the small trapping area and lack of need to return to the primary oviposition site of the village. Notably, the average displacement distance of recaptured mosquitoes was 136 m, and the maximum was 623 m (Fig. 4), whereas the average distance between all release points and all houses of the village was 230 m (maximum=886 m). The difference implies that mosquito movement is not random and probably reflects preference sites that are richer in resources (larval sites, host-feeding sites, etc.), which were concentrated in the center zones (Fig. 1). High agreement between Monte Carlo simulated distances of mosquitoes moving randomly from the released zones to all houses (r^2^=94%, P=0.006, Fig. 4d) with a slope of 1 m/m provides strong support that the layout and size of the trapping area had a major effect on the movement pattern. Moreover, it implies that displacement distances in this experiment was depended on the layout to the houses (traps). Greatly limiting inference on the behavior of these mosquitoes including their movement distance kernel. The negative intercept (-102 m) suggests that the most extreme houses, in the south, east, and north zones were less likely to be visited by mosquitoes, albeit this in part may be due to small number of recaptures (Figs. 1 and 4). In our study the areas with highest density of houses (center zones) were closest to the main larval site, preventing a separation of the specific effects of these factors.

Mosquitoes exhibit multiple modalities of movements. Typical appetitive movements aimed at seeking nearby resources, e.g., resting sites, hosts, sugar sources, mates, and larval sites occur within the flight boundary layer (<6 m above ground) where the mosquito’s own flight speed and direction determines its direction and displacement. Because on average flight speed does not exceed 1 m/s (Clements 1955), the displacement associated with appetitive movements span a small scale of (<10 km). Additionally, mosquitoes engage in high-altitude (100-300 m above ground) windborne migration (Glick 1939, Reynolds et al. 1996, Kay and Farrow 2000, Huestis et al. 2019, Sanogo et al. 2021, Yaro et al. 2022, Atieli et al. 2023, Bamou et al. 2025), as many other insects (Drake and Reynolds 2012, Chapman et al. 2015, Florio et al. 2020, Nartey et al. 2024, Sappington 2024). Taking advantage of fast winds (4-12 m/s) at altitude, they cover tens if not hundreds of kilometers in a single night (Sellers et al. 1977, 1982, Pedgley 1983, Sellers 1989, Sellers and Maarouf 1990, Huestis et al. 2019). While all mosquitoes engage in appetitive movements, it is not known what fraction of the population engages in high-altitude windborne flights. However, given that inclusive daily mortality (emigration included) is 5-20% in *A. gambiae* s.l. (Guerra et al. 2014, Matthews et al. 2020, Matthews 2025), the fraction in this species group cannot be expected to exceed 5–10%, except during short periods with specific conditions, which remain unknown. The current study aimed at understanding the first modality of movement that includes appetitive flights.

Although our MRR experiment lasted 101 days, our marking method allowed detection of recaptures up to 36 days post release (minimum age). One mosquito was recaptured 30 d after release and two were caught after 21 d. The probability of daily survival (PDS) was 86% (Gillies) and 94% after correction (Buonaccorsi et al. 2003), which are at the higher end of the MRR estimates. The length of the recapture period had a strong effect on the estimate of daily survival (PDS) as illustrated in the inset of Fig. 4d. Using the widely used regression method to estimate PDS, the estimate based on the 12 days recapture is 0.78 and is systematically increasing to 0.86 for 30 days. While we cannot be sure that there no mosquitoes with minimum age greater than 36 days (Methods), these results demonstrate that many studies that used short period of recapture (<12 d, above) likely underestimated daily survival.

The age composition of the mosquitoes of Thierola has dramatically changed over the course of the experiment, from the very young population in September (peak density) reflecting large influx of emergent adults, the increase in population age as this influx (and density) declines, to the old population during the minimum (mid-November–early December, Fig. 2b and inset). Although not statistically significant due to the small sample size, infection rate among females that died during the handling and marking (3.3% overall) increased from 2.7% (September N=184) to 7.7% (October– December, N=26), consistent with expectations based on older population.

The overall sporozoites infection rate in recaptured mosquitoes was 23.9% (N=46). This rate itself reflects the aging population of the mosquitoes towards the later part of the wet season and the onset of the dry season (above). Considering the “cohort” of captured and released mosquitoes, the sporozoite rate increased 15% per gonotrophic cycle (3 d, Fig. S2). This rate of increase considers only surviving mosquitoes, so mortality is not part of this rate. If all mosquitoes that bite an infectious person develop sporozoites without change in mortality, this rate also estimates the fraction of bloodmeals on infectious persons in Thierola at this period. Because estimates of gametocyte carriers Sahelian communities are typically higher than 15% (Ouedraogo et al. 2010), it is likely that at least one of the above assumptions is incorrect.

### Lessons for improving estimation of survival, movement distances, and population sizes

Our results highlight that most MRR experiments on *Anopheles* mosquitoes had too small trapping areas and were carried out over too short duration, thus underestimated daily survival, movement distances, and population size, echoing concerns that were raised in meta-analyses (Guerra et al. 2014). Indeed, given that movement measured in our MRR was heavily shaped by the small trapping area layout, we could not derive meaningful movement statistics accept for testing site fidelity. Accordingly, effective experimental design ensures spatial and temporal margins demonstrating the limits of movement and longevity. We propose 300-600 m wide outer “ring” of traps without recaptures, surrounding the furthest distance reached by any recaptured mosquitoes. This ring should have equal (or sufficient) trapping effort to avoid confounding lack of recaptures with lack of effort to find them. Likewise, the duration of recapture collections should ensure that the last 3-6 capture dates collect no recaptured mosquitoes (despite sufficient trapping effort). Such experiments are considerably larger and accordingly require greater resources in terms of trained entomologists, collectors, funding, and deploy a marking approach that facilitates discrimination between multiple groups (Faiman et al. 2021). Collection interval matching the length of the gonotrophic cycle of the target species at the time of the experiment (typically 3 d for anophelines in tropical areas) serves to focus on net displacement by averaging out movements that aimed at different resources in multiple and often opposing directions, e.g., larval sites and hosts concentration. Despite higher cost, such experiments will pay off by resolving inconsistencies and yielding useful parameters to improve understanding of disease transmission and accurately model disease burden and control campaigns.

### Ethical considerations

This protocol has been submitted to the USTTB ethics committee for approval under No. 2019/82/CE/FMPOS. At the start of activities, the study protocol and objectives were clearly explained to the local authorities. After obtaining community consent from the local chiefdom, individual consent was sought and obtained from all participants. Throughout the study period, the following ethical aspects were respected: the collaboration of each villager was free and voluntary; access to the houses for mosquito catching was conditional on the favorable opinion of the head of the family and the owner. The guides and collectors were people from local communities, well known to each other and chosen by the heads of the families in each village.

## Acknowledgements

We are grateful to the residents of Thierola and local authorities for their cooperation and support and hospitality. We are indebted to the field and laboratory teams for their dedication and technical expertise in mosquito collection, marking, and analysis. We thank Ms. Margory Sullivan, Ms. Fatoumata Bathili, Mr. Samuel Moretz, Drs. Thomas Wellems (National Institutes of Health, USA) for crucial logistical support; Drs. We thank Drs. Tom Burkot (Australian Institute of Tropical Health and Medicine), Hamidou Maiga (IRSS, Burkina Faso), Nafomon Sogoba, Mahamadou B. Touré, Amadou Sekou Traoré, and Moussa Keita for their valuable feedback on earlier drafts of this manuscript.

## Supplemental Information

**Figure S1.**
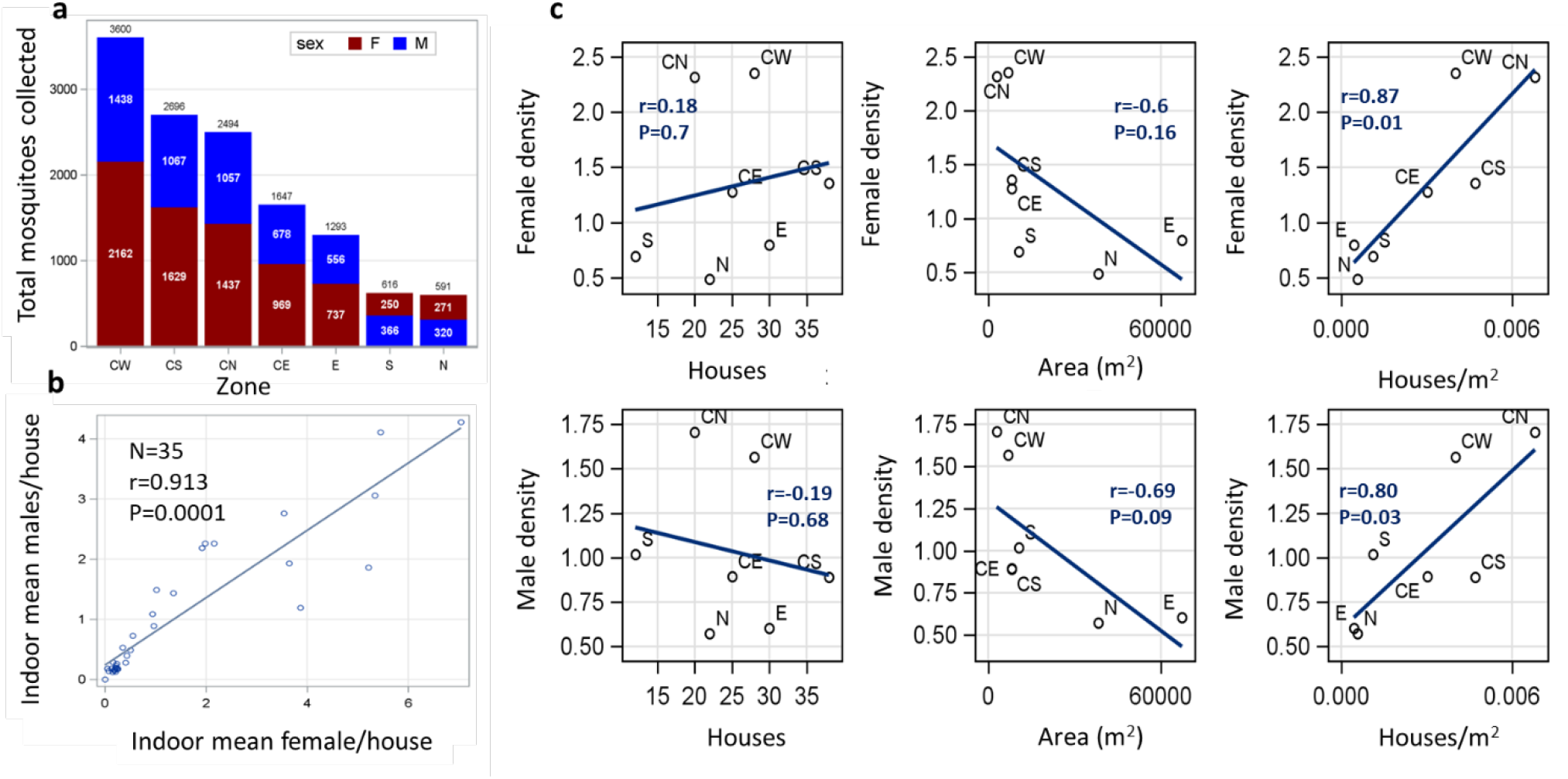
Mosquito capture variation between zones and species. The total number of females and males collected per zone (a). The relationship between mean indoor density of males and females in each day of collection with Pearson Correlation coefficient (b). The relationships of female and male indoor density per zone to the number of houses per zone, the approximate area of each zone, and the house density per zone (c). Trend (regression) lines and correlation coefficients are shown.

**Figure S2.**
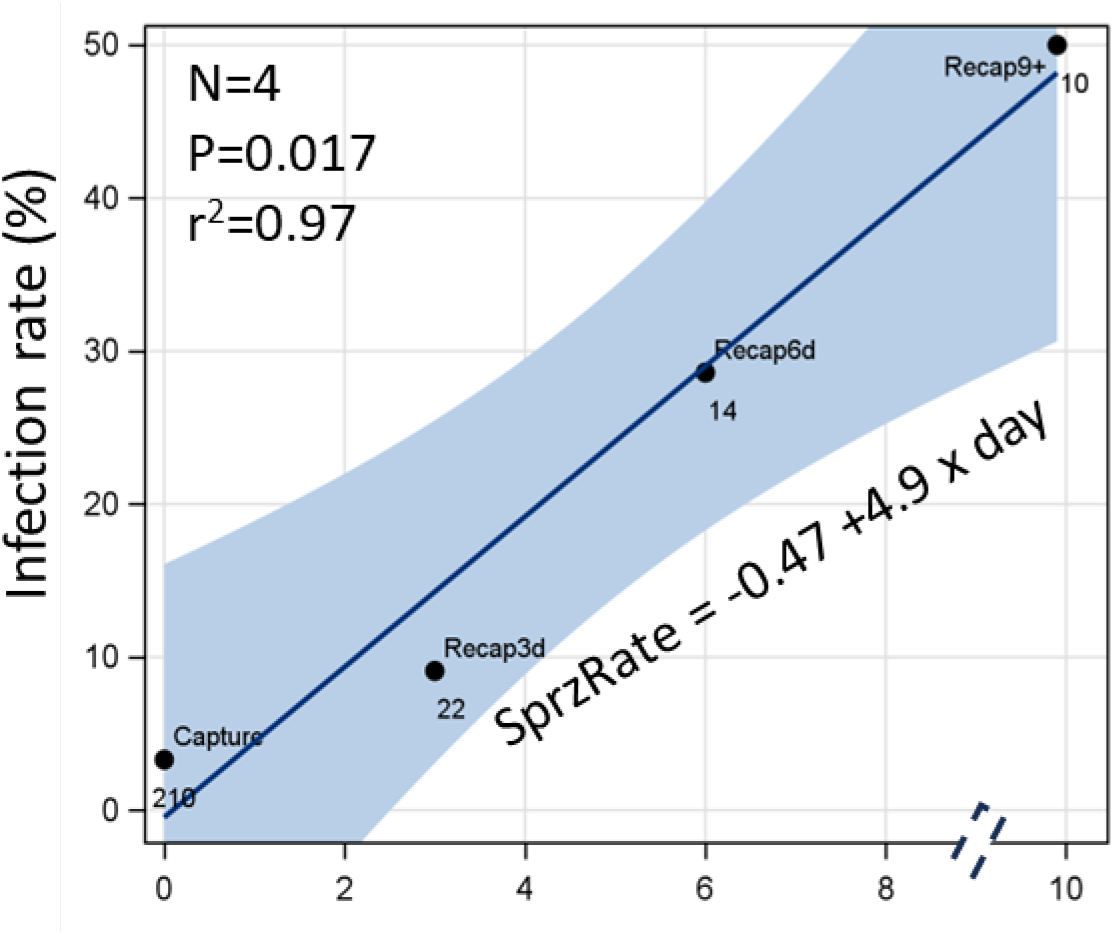
Increase in sporozoite infection rate (thorax-head infection) among recaptured females over time since release. Sample size per time point is given below each point and its time period above. Note that due to small sample size, recaptured females collected 9 or more days after release were pooled, denoted as ‘Recap9+’ and broken X axis. Time zero relates to the sporozoite infection rate measured at capture among dead females. Linear regression and 95%CI of the expected mean are shown with regression equation, statistical significance (P).

## References

Adamou A, Dao A, Timbine S et al. 2011. The contribution of aestivating mosquitoes to the persistence of Anopheles gambiae in the Sahel. Malar J 10(151. 10.1186/1475-2875-10-151.

Alonso WJ, Schuck-Paim C. 2006. The ‘ghosts’ that pester studies on learning in mosquitoes: guidelines to chase them off. Med Vet Entomol 20(2):157–165. 10.1111/j.1365-2915.2006.00623.x.

Atieli HE, Zhou G, Zhong D et al. 2023. Wind-assisted high-altitude dispersal of mosquitoes and other insects in East Africa. J Med Entomol 60(4):698–707. 10.1093/jme/tjad033.

Bamou R, Dao A, Yaro AS et al. 2025. Pathogens spread by high-flying wind-borne mosquitoes. Proc Natl Acad Sci U S A 122(48):e2513739122. 10.1073/pnas.2513739122.

Brady OJ, Godfray HC, Tatem AJ et al. 2016. Vectorial capacity and vector control: reconsidering sensitivity to parameters for malaria elimination. Trans R Soc Trop Med Hyg 110(2):107–117. 10.1093/trstmh/trv113.

Buonaccorsi JP, Harrington LC, Edman JD. 2003. Estimation and comparison of mosquito survival rates with release-recapture-removal data. J Med Entomol 40(1):6–17. 10.1603/0022-2585-40.1.6.

Carter R, Mendis KN, Roberts D. 2000. Spatial targeting of interventions against malaria. Bull World Health Organ 78(12):1401–1411.

Chapman JW, Reynolds DR, Wilson K. 2015. Long-range seasonal migration in insects: mechanisms, evolutionary drivers and ecological consequences. Ecology Letters 18(3):287–302. 10.1111/ele.12407.

Charlwood JD, Graves PM, Marshall TF. 1988. Evidence for a ‘memorized’ home range in Anopheles farauti females from Papua New Guinea. Med Vet Entomol 2(2):101–108. 10.1111/j.1365-2915.1988.tb00059.x.

Chitnis N, Hyman JM, Cushing JM. 2008. Determining important parameters in the spread of malaria through the sensitivity analysis of a mathematical model. Bull Math Biol 70(5):1272–1296. 10.1007/s11538-008-9299-0.

Clements AN. 1955. The Sources of Energy for Flight in Mosquitoes. Journal of Experimental Biology 32(3):547–554.

Costantini C, Li SG, DellaTorre A et al. 1996. Density, survival and dispersal of Anopheles gambiae complex mosquitoes in a West African Sudan savanna village. Medical and Veterinary Entomology 10(3):203–219. 10.1111/j.1365-2915.1996.tb00733.x.

Dao A, Yaro AS, Diallo M et al. 2014. Signatures of aestivation and migration in Sahelian malaria mosquito populations. Nature 516(7531):387–390. 10.1038/nature13987.

Drake VA, Reynolds DR. 2012. Radar Entomology: Observing Insect Flight and Migration. CABI International.

Faiman R, Yaro AS, Dao A et al. 2022. Isotopic evidence that aestivation allows malaria mosquitoes to persist through the dry season in the Sahel. Nat Ecol Evol 6(11):1687–1699. 10.1038/s41559-022-01886-w.

Faiman R, Krajacich BJ, Graber L et al. 2021a. A novel fluorescence and DNA combination for versatile, long-term marking of mosquitoes. Methods Ecol Evol 12(6):1008–1016. 10.1111/2041-210X.13592.

Faiman R, Krajacich BJ, Graber L et al. 2021b. A novel fluorescence and DNA combination for versatile, long-term marking of mosquitoes. Methods in Ecology and Evolution 12(6):1008–1016. 10.1111/2041-210x.13592.

Florio J, Verú LM, Dao A et al. 2020. Diversity, dynamics, direction, and magnitude of high-altitude migrating insects in the Sahel. Sci Rep-Uk 10(1). https://doi.org/ARTN p20523 10.1038/s41598-020-77196-7.

Gillies MT. 1961. Studies on the dispersion and survival of Anopheles gambiae Giles in East Africa, by means of marking and release experiments. B Entomol Res 52(1):99–127. 10.1017/S0007485300055309.

Gillies MT, Wilkes TJ. 1965. A Study of Age-Composition of Populations of Anopheles Gambiae Giles and a Funestus Giles in North-Eastern Tanzania. B Entomol Res 56(237–+. 10.1017/S0007485300056339.

Glick PA. 1939. The Distribution of Insects, Spiders and Mites in the Air

Guerra CA, Reiner RC, Perkins TA et al. 2014. A global assembly of adult female mosquito mark-release-recapture data to inform the control of mosquito-borne pathogens. Parasite Vector 7(https://doi.org/Artn p276 10.1186/1756-3305-7-276.

Huestis DL, Dao A, Diallo M et al. 2019. Windborne long-distance migration of malaria mosquitoes in the Sahel. Nature 574(7778):404–408. 10.1038/s41586-019-1622-4.

Kamdem C, Fouet C, Etouna J et al. 2012. Spatially Explicit Analyses of Anopheline Mosquitoes Indoor Resting Density: Implications for Malaria Control. Plos One 7(2). https://doi.org/ARTN pe31843 10.1371/journal.pone.0031843.

Kay BH, Farrow RA. 2000. Mosquito (Diptera: Culicidae) dispersal: implications for the epidemiology of Japanese and Murray Valley encephalitis viruses in Australia. J Med Entomol 37(6):797–801. 10.1603/0022-2585-37.6.797.

Lehmann T, Dao A, Yaro AS et al. 2010. Aestivation of the African malaria mosquito, Anopheles gambiae in the Sahel. Am J Trop Med Hyg 83(3):601–606. 10.4269/ajtmh.2010.09-0779.

Lehmann T, Dao A, Yaro AS et al. 2014. Seasonal variation in spatial distributions of Anopheles gambiae in a Sahelian village: evidence for aestivation. J Med Entomol 51(1):27–38. 10.1603/me13094.

Matthews J. 2025. Mosquito survival from mark-recapture studies releasing at known age. Parasite Vector 18(1). https://doi.org/ARTN p455 10.1186/s13071-025-07024-2.

Matthews J, Bethel A, Osei G. 2020. An overview of malarial Anopheles mosquito survival estimates in relation to methodology. Parasit Vectors 13(1):233. 10.1186/s13071-020-04092-4.

McCall PJ, Kelly DW. 2002. Learning and memory in disease vectors. Trends in Parasitology 18(10):429–433. https://doi.org/PiiS1471-4922(02)02370-X Doi 10.1016/S1471-4922(02)02370-X.

McCall PJ, Mosha FW, Njunwa KJ et al. 2001. Evidence for memorized site-fidelity in. T Roy Soc Trop Med H 95(6):587–590. 10.1016/S0035-9203(01)90087-2.

Nartey R, Chamorro ML, Buffington M et al. 2024. Invasion and spread of the neotropical leafhopper Curtara insularis (Hemiptera, Cicadellidae) in Africa and North America and the role of high-altitude windborne migration in invasive insects. Neobiota 96(173–189.

Omer SM, Cloudsley-Thompson JL. 1970. Survival of female Anopheles gambiae Giles through a 9-month dry season in Sudan. Bull World Health Organ 42(2):319–330.

Ouedraogo AL, Bousema T, de Vlas SJ et al. 2010. The plasticity of Plasmodium falciparum gametocytaemia in relation to age in Burkina Faso. Malar J 9(281. 10.1186/1475-2875-9-281.

Pedgley DE. 1983. Windborne Spread of Insect-Transmitted Diseases of Animals and Man. Philos T R Soc B 302(1111):463–470. 10.1098/rstb.1983.0068.

Reisen WK, Milby MM, Meyer RP et al. 1991. Mark Release Recapture Studies with Culex Mosquitos (Diptera, Culicidae) in Southern California. Journal of Medical Entomology 28(3):357–371. 10.1093/jmedent/28.3.357.

Reynolds DR, Smith AD, Mukhopadhyay S et al. 1996. Atmospheric transport of mosquitoes in northeast India. Medical and Veterinary Entomology 10(2):185–186. 10.1111/j.1365-2915.1996.tb00727.x.

Ribeiro JMC, Seulu F, Abose T et al. 1996. Temporal and spatial distribution of anopheline mosquitos in an Ethiopian village: Implications for malaria control strategies. B World Health Organ 74(3):299–305.

Sanogo ZL, Yaro AS, Dao A et al. 2021. The Effects of High-Altitude Windborne Migration on Survival, Oviposition, and Blood-Feeding of the African Malaria Mosquito, Anopheles gambiae s.l. (Diptera: Culicidae). J Med Entomol 58(1):343–349. 10.1093/jme/tjaa137.

Sappington TW. 2024. Aseasonal, undirected migration in insects: ‘Invisible’ but common. iScience 27(6):110040. 10.1016/j.isci.2024.110040.

SAS software. 2019. ‘Chapter SAS Institute Inc’. Cary, NC, USA.

Sellers RF. 1989. Eastern equine encephalitis in Quebec and Connecticut, 1972: introduction by infected mosquitoes on the wind? Can J Vet Res 53(1):76–79.

Sellers RF, Maarouf AR. 1990. Trajectory analysis of winds and eastern equine encephalitis in USA, 1980-5. Epidemiol Infect 104(2):329–343. 10.1017/s0950268800059501.

Sellers RF, Pedgley DE, Tucker MR. 1977. Possible Spread of African Horse Sickness on Wind. J Hyg-Cambridge 79(2):279–298. 10.1017/S0022172400053109.

Sellers RF, Pedgley DE, Tucker MR. 1982. Rift Valley fever, Egypt 1977: disease spread by windborne insect vectors? Vet Rec 110(4):73–77. 10.1136/vr.110.4.73.

Service MW. 1993. Mosquito Ecology: Field Sampling Methods, 2nd. London: Elsevier Applied Science.

Smith T, Charlwood JD, Takken W et al. 1995. Mapping the Densities of Malaria Vectors within a Single Village. Acta Tropica 59(1):1–18. 10.1016/0001-706x(94)00082-C.

Toure YT, Dolo G, Petrarca V et al. 1998. Mark-release-recapture experiments with Anopheles gambiae sl in Banambani Village, Mali, to determine population size and structure. Medical and Veterinary Entomology 12(1):74–83. 10.1046/j.1365-2915.1998.00071.x.

Verdonschot PFM, Besse-Lototskaya AA. 2014. Flight distance of mosquitoes (Culicidae): A metadata analysis to support the management of barrier zones around rewetted and newly constructed wetlands. Limnologica 45(69–79. 10.1016/j.limno.2013.11.002.

WHO. 2025. World malaria report: addressing the threat of antimalarial drug resistance. Available from: https://www.who.int/teams/global-malaria-programme/reports/world-malaria-report-2025

Yaro AS, Linton YM, Dao A et al. 2022. Diversity, composition, altitude, and seasonality of high-altitude windborne migrating mosquitoes in the Sahel: Implications for disease transmission. Frontiers in Epidemiology 13 October 2022 (10.3389/fepid.2022.1001782.

